# *massDatabase*: utilities for the operation of the public compound and pathway database

**DOI:** 10.1101/2022.06.02.494457

**Authors:** Xiaotao Shen, Chuchu Wang, Michael P. Snyder

**Author notes:** These authors contributed equally (Xiaotao Shen, Chuchu Wang).

## Abstract

**Summary:** One of the major challenges in LC-MS data (metabolome, lipidome, and exposome) is converting many metabolic feature entries to biological function information, such as metabolite annotation and pathway enrichment, which are based on the compound and pathway databases. Multiple online databases have been developed, containing lots of information about compounds and pathways. However, there is still no tool developed for operating all these databases for biological analysis. Therefore, we developed *massDatabase*, an R package that operates the online public databases and combines with other tools for streamlined compound annotation and pathway enrichment analysis. *massDatabase* is a flexible, simple, and powerful tool that can be installed on all platforms, allowing the users to leverage all the online public databases for biological function mining. A detailed tutorial and a case study are provided in the **Supplementary Materials**.

**Availability and implementation:** https://massdatabase.tidymass.org/.

**Supplementary information:** Supplementary data are available at *Bioinformatics* online.

## 1 Introduction

Liquid chromatography coupled to mass spectrometry (LC−MS) is a comprehensive, unbiased technology to research small compounds, which has become increasingly popular in dietary, environmental, and biomedical studies (Wishart, 2016). One of the major challenges in LC-MS data (metabolome, lipidome, and exposome) is the post-processing of a large number of metabolic feature entries to achieve clear biological evidence, such as the compound annotation and pathway enrichment. Therefore, the databases for compounds and pathways are essential for these analyses. Multiple public databases for compounds and pathways are available online, which benefits the community (Go, 2010). However, the existence of an automated, multiple compound/pathway query processing package in R is still a demand. So far, although several R packages have been developed to extract online databases, most of them only support one or limited databases and have different design concepts and output formats. In addition, they can not be combined with other existing tools for a straightforward subsequent analysis, which limits their further applications.

Here, we presented the massDatabase package to overcome the challenges mentioned above while accessing the online databases, particularly to (1) support most of the commonly used online public databases (11 databases, **Table S1**), (2) operate (extracting, downloading, reading, and converting) the online public databases, and (3) combine the online public databases with existing tools for subsequent compound annotation, and pathway enrichment analysis (**Fig. 1**).

**Fig. 1.**
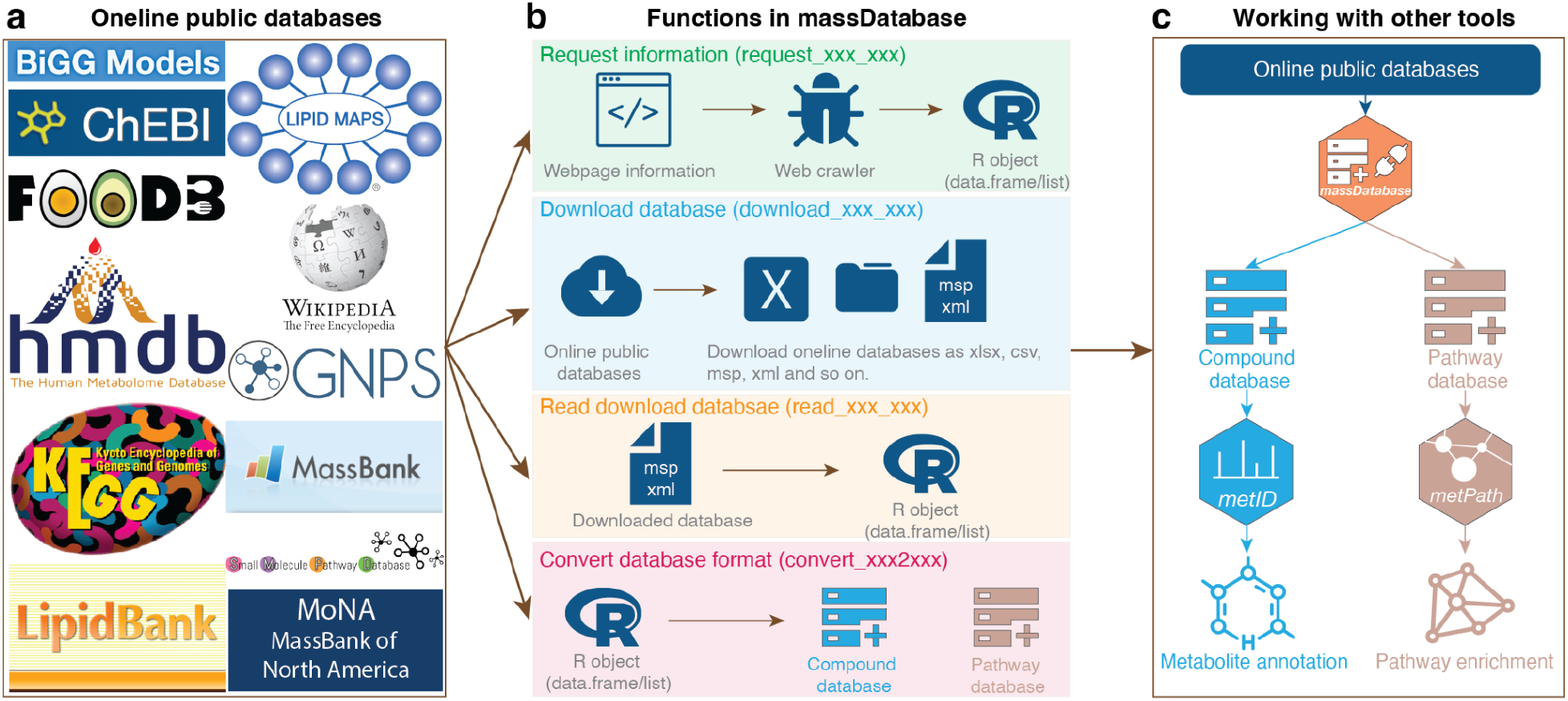
The overview of **(a)** the online databases that massDatabase support, **(b)** the functions used to process databases, and **(c)** the combination with other tools in the tidyMass project.

## 2 Features and methods

Using *massDatabase*, users can extract compound/pathway information from the online databases (11 databases, **Table S1**) and download them. In addition, *massDatabase* can also be combined with other tools for metabolite annotation and pathway enrichment analysis. The *massDatabase* can be installed on Mac OS, Windows, and Linux.

### 2.1 Online databases operation

The functions in *massDatabase* could be grouped into four classes. (1) Request specific information of one item (compound, pathway, reaction, *etc*.) online using the web crawler, (2) download the corresponding database, (3) read the downloaded databases (csv, mgf formats, *etc*.) as R object (list or data frame), and (4) convert the databases to other formats that could be used for other tools (**Fig. 1**).

### 2.2 Combination with other tools

The users can download the online databases and then convert them to the formats supported by the packages in the tidyMass project using *massDatabase*. Currently, two packages from tidyMass projects could combine with *massDatabase*. Users can download the compound databases (MS^1^ or MS^2^ spectra databases), convert them to the database format in the *metID* package, and then use them for compound annotation by *metID*. Furthermore, users can also download the pathway databases, convert them to the pathway database format in the *metPath* package, and then use them for pathway enrichment analysis by *metPath*.

## 3 Case study

We applied *massDatabase* to a published study from our lab (Liang *et al*., 2020) as a case study for exemplifying the value of *massDatabase* in biological function mining by integrating with the online public databases. The MS^2^ spectra databases from HMDB, MassBank, and MoNA were first downloaded and converted to databases format in *metID*. And the pathway database from KEGG is downloaded and converted to pathway database format in *metPath*. Then the metabolic feature table was annotated by *metID* which is based on the public databases from *massDatabase* and our in-house library. Then all the annotated metabolites were used for pathway enrichment analysis using *metPath*. The top enriched pathways include Steroid hormone biosynthesis, Phenylalanine metabolism, Caffeine metabolism, Linoleic acid metabolism, Primary bile acid biosynthesis, *etc*., which are most consistent with the original analysis (**Fig. S1**) (Liang *et al*., 2020). These results indicate that *massDatabase* is a powerful tool for utilizing online public compound and pathway databases for automated and reproducible analysis of LC-MS-based metabolomics data (**Supplementary Material**).

## 4 Conclusion

*massDatabase* is developed to operate public databases in untargeted LC-MS-based data (metabolome, lipidome, and exosome). It allows users to extract, download, read databases, and convert database formats to different formats required by other tools. To our best knowledge, it is the first R package allowing users to operate most of the commonly used online public databases for subsequent biological function mining.

