## Supplementary information for "*massDatabase*: utilities for the operation of the public compound and pathway database"

[+] These authors contributed equally.
[*] To whom correspondence should be addressed.

**Table S1. The online public databases are supported by *massDatabase*.**

| **Database** | **Type** | $MS^2$ spectra for compounds | Link |
|---|---|---|---|
| BiGG model | Compounds and reactions | No | http://bigg.ucsd.edu/ |
| ChEBI | Compounds | No | https://www.ebi.ac.uk/chebi/ |
| FoodB | Compounds | No | https://foodb.ca/ |
| HMDB | Metabolites | Yes | https://hmdb.ca/ |
| KEGG | Metabolites, drugs, reaction and pathays | No | https://www.genome.jp/kegg |
| LipidBank | Lipids | No | https://lipidbank.jp/ |
| LipidMaps | Lipids | No | https://www.lipidmaps.org/ |
| GNPS | Compounds | Yes | https://gnps.ucsd.edu/ProteoSAFe/static/gnps-splash.jsp |
| MassBank | Compounds | Yes | https://massbank.eu/ |
| SMPDB | Pathways | - | https://www.smpdb.ca/ |
| MoNA | Compounds | Yes | https://mona.fiehnlab.ucdavis.edu/ |

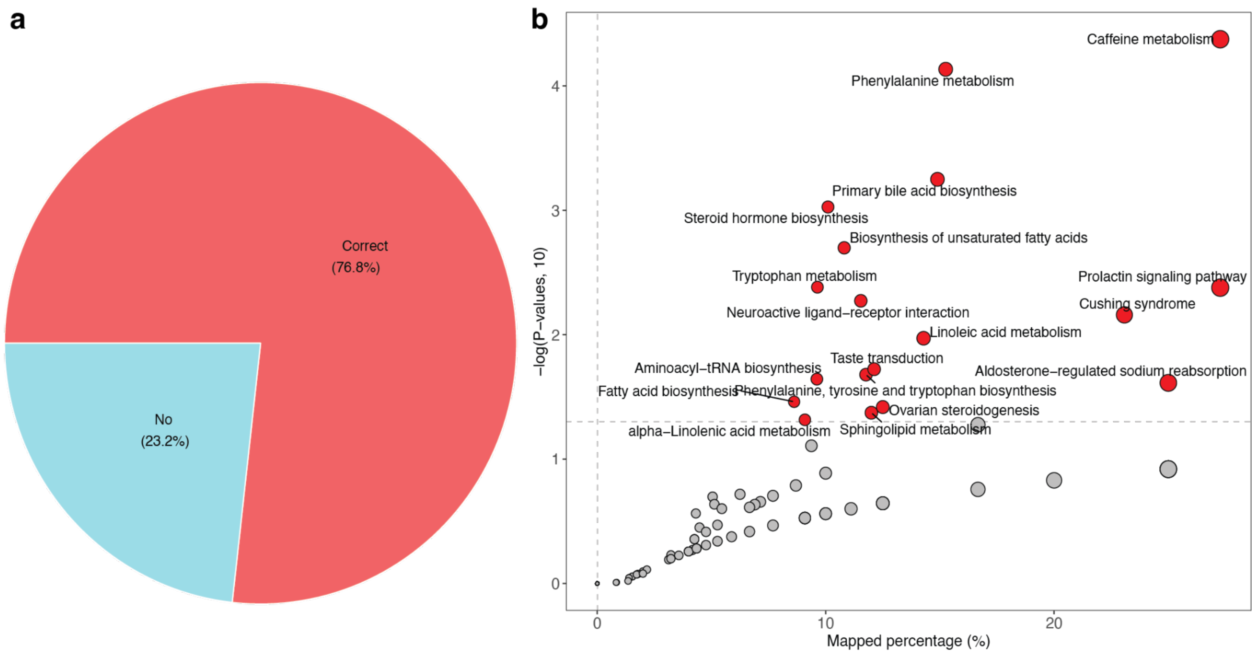


**Figure S1. Analysis result of case study using *metID* and *metPath*. (a)** The annotation result from HMDB, MassBank and MoNA compared to annotation results from in-house databases. (**b**) Pathway enrichment analysis results.

### Supplementary Note

The code used to download the HMDB, MoNA and MassBank compound databases, and KEGG pathway databases. The code of *massDatabase* could be found on GitHub (https://github.com/tidymass/massdatabase/).

### 1. HMDB database

```
######
no_source()

masstools::setwd_project()
rm(list = ls())
load("other_files/HMDB/MS1/hmdb_ms1.rda")
setwd("other_files/HMDB/MS2/")

library(tidyverse)
library(xml2)
library(stringr)

# file <-
# dir("hmdb_experimental_msms_spectra/")
#
# hmdb_ms2 <-
#   file %>%
# purrr::map(function(x){
#   cat(x, " ")
```

```
#   data <-
# read_xml(file.path("hmdb_experimental_msms_spectra/", x)) %>%
#     xml2::as_list()
#   Instrument_type <-
#   unlist(data$`ms-ms`$`instrument-type`)
#   Instrument_type <-
#     ifelse(is.null(Instrument_type), NA, Instrument_type)
#   Polarity <-
#     unlist(data$`ms-ms`$`ionization-mode`)
#   Polarity <-
#     ifelse(is.null(Polarity), NA, Polarity)
#   collision_energy_level <-
#   unlist(data$`ms-ms`$`collision-energy-level`)
#   collision_energy_level <-
#     ifelse(is.null(collision_energy_level), NA, collision_energy_level)
#   collision_energy_voltage <-
#     unlist(data$`ms-ms`$`collision-energy-voltage`)
#   collision_energy_voltage <-
#     ifelse(is.null(collision_energy_voltage), NA, collision_energy_voltage)
#
#   chromatography_type <-
#     unlist(data$`ms-ms`$`chromatography-type`)
#   chromatography_type <-
#     ifelse(is.null(chromatography_type), NA, chromatography_type)
#   analyzer_type <-
#     unlist(data$`ms-ms`$`analyzer-type`)
#   analyzer_type <-
#     ifelse(is.null(analyzer_type), NA, analyzer_type)
#   ionization_type <-
#     unlist(data$`ms-ms`$`ionization-type`)
#   ionization_type <-
#     ifelse(is.null(ionization_type), NA, ionization_type)
#   charge_type <-
#     unlist(data$`ms-ms`$`charge-type`)
#   charge_type <-
#     ifelse(is.null(charge_type), NA, charge_type)
#   adduct <-
#     unlist(data$`ms-ms`$adduct)
#   adduct <-
#     ifelse(is.null(adduct), NA, adduct)
#   adduct_type <-
#     unlist(data$`ms-ms`$`adduct-type`)
#   adduct_type <-
#     ifelse(is.null(adduct_type), NA, adduct_type)
#   adduct_mass <-
#     unlist(data$`ms-ms`$`adduct-mass`)
#   adduct_mass <-
#     ifelse(is.null(adduct_mass), NA, adduct_mass)
#   ms1_info <-
#     data.frame(HMDB.ID = stringr::str_extract(x, "HMDB[0-9]{7,9}"),
```

```
#               Instrument_type = Instrument_type,
#               Polarity = Polarity, collision_energy_level = collision_energy_level,
#               collision_energy_voltage = collision_energy_voltage,
#               chromatography_type = chromatography_type,
#               analyzer_type = analyzer_type,
#               ionization_type = ionization_type,
#               charge_type = charge_type,
#               adduct = adduct,
#               adduct_type = adduct_type,
#               adduct_mass = adduct_mass)
#
#   ms2 <-
#     data$`ms-ms`$`ms-ms-peaks`
#   if(is.null(ms2)){
#     ms2 <- data.frame()
#   }else{
#     ms2 <-
#       lapply(ms2, function(y){
#         unlist(y)
#       }) %>%
#       dplyr::bind_rows() %>%
#       as.data.frame() %>%
#       dplyr::select(`mass-charge`, intensity) %>%
#       dplyr::rename(mz = `mass-charge`)
#   }
#   list(ms1_info = ms1_info,
#        ms2 = ms2)
# })
#
# remove_idx <-
# hmdb_ms2 %>%
#   lapply(function(x) {
#     nrow(x$ms2)
#   }) %>%
#   unlist() %>%
#   `==`(0) %>%
#   which()
#
# hmdb_ms2[[16776]]$ms2
#
# hmdb_ms2 <-
#   hmdb_ms2[-16776]
#
#
# spectra_info <-
#   hmdb_ms2 %>%
#   purrr::map(function(x){
#     x$ms1_info
#   }) %>%
#   dplyr::bind_rows() %>%
```

```
#   as.data.frame()
#
# save(spectra_info, file = "spectra_info")
#
# spectra_data <-
#   hmdb_ms2 %>%
#   purrr::map(function(x){
#     x$ms2
#   })
#
# save(spectra_data, file = "spectra_data")

load("spectra_info")
load("spectra_data")

spectra_info$HMDB.ID

spectra_info[which(spectra_info == "NA", arr.ind = TRUE)] <- NA
spectra_info[which(spectra_info == "n/a", arr.ind = TRUE)] <- NA
spectra_info[which(spectra_info == "N/A", arr.ind = TRUE)] <- NA

spectra_info <-
spectra_info %>%
  dplyr::select(HMDB.ID, Instrument_type, Polarity, collision_energy_voltage, adduct)

spectra_info$Polarity

remove_idx <-
  which(is.na(spectra_info$Polarity))
remove_idx
dim(spectra_info)
length(spectra_data)

spectra_info <-
  spectra_info[-remove_idx,]

spectra_data <-
  spectra_data[-remove_idx]

spectra_info <-
  spectra_info %>%
  dplyr::mutate(Polarity = case_when(
    Polarity == "positive" ~ "Positive",
    Polarity == "negative" ~ "Negative",
    TRUE ~ Polarity
  ))

library(plyr)

spectra_info$Lab.ID <-
```

```
  masstools::name_duplicated(spectra_info$HMDB.ID) %>%
  paste("shen", sep = "_")

spectra_info2 <-
  spectra_info %>%
  plyr::dlply(.variables = .(HMDB.ID)) %>%
  purrr::map(function(y) {
    if (sum(is.na(y$collision_energy_voltage)) > 0) {
      y$collision_energy_voltage[is.na(y$collision_energy_voltage)] <-
        paste("Unknown",
1:length(y$collision_energy_voltage[is.na(y$collision_energy_voltage)]), sep = "_")
    }
    y
  }) %>%
  dplyr::bind_rows() %>%
  as.data.frame()

spectra_info2 <-
  spectra_info2[match(spectra_info$Lab.ID, spectra_info2$Lab.ID),]

sum(spectra_info2$Lab.ID == spectra_info$Lab.ID)

spectra_data2 <-
1:length(spectra_data) %>%
  purrr::map(function(i){
    x <- spectra_data[[i]]
    x <- list(x)
    names(x) <-
      spectra_info2$collision_energy_voltage[i]
    x
  })

names(spectra_data2) <- spectra_info2$Lab.ID

######positive mode
spectra_info2$Lab.ID == names(spectra_data2)

index_pos <- which(spectra_info2$Polarity == "Positive")
index_neg <- which(spectra_info2$Polarity == "Negative")

spectra_info_pos <- spectra_info2[index_pos,]
spectra_data_pos <- spectra_data2[index_pos]

spectra_info_neg <- spectra_info2[index_neg,]
spectra_data_neg <- spectra_data2[index_neg]

colnames(spectra_info2)
colnames

spectra_info2 <-
```

```
spectra_info2 %>%
  dplyr::rename(CE = "collision_energy_voltage")

spectra_info2 <-
spectra_info2 %>%
  dplyr::left_join( %>% dplyr::select(-Lab.ID), by = c("HMDB.ID"))

spectra_info2$mz

hmdb_ms2 <- hmdb_ms1

hmdb_ms2@spectra.info <- spectra_info2

hmdb_ms2@spectra.data$Spectra.positive <- spectra_data_pos
hmdb_ms2@spectra.data$Spectra.negative <- spectra_data_neg

hmdb_ms2

 %>%
  dplyr::count(HMDB.ID, Polarity)

idx <-
 %>%
dplyr::filter(HMDB.ID == "HMDB0000288") %>%
  pull(Lab.ID)

idx2 <-
which(names(hmdb_ms2@spectra.data$Spectra.positive) %in% idx)

masstools::ms2_plot(spectrum1 = hmdb_ms2@spectra.data$Spectra.positive[[idx2[1]]][[1]],
                    spectrum2 = hmdb_ms2@spectra.data$Spectra.positive[[idx2[5]]][[1]])


head(hmdb_ms2@spectra.info$mz)
head(hmdb_ms2@spectra.info$monisotopic_molecular_weight)

hmdb_ms2@spectra.data$Spectra.positive <-
  hmdb_ms2@spectra.data$Spectra.positive %>%
  purrr::map(function(x){
    x %>%
      lapply(function(y){
        y$mz <- as.numeric(y$mz)
        y$intensity <- as.numeric(y$intensity)
        y
      })
  })

hmdb_ms2@spectra.data$Spectra.negative <-
  hmdb_ms2@spectra.data$Spectra.negative %>%
  purrr::map(function(x){
```

```
    x %>%
      lapply(function(y){
        y$mz <- as.numeric(y$mz)
        y$intensity <- as.numeric(y$intensity)
        y
      })
  })

save(hmdb_ms2, file = "hmdb_ms2.rda")
```

**2. MoNA database**

```
###source
no_source()

masstools::setwd_project()
rm(list = ls())
load("other_files/HMDB/MS1/hmdb_ms1.rda")
load("other_files/KEGG/kegg_ms1.rda")
source("R/read_msp_data.R")
source("R/3_utils.R")
source("R/10_MASSBANK.R")
setwd("other_files/MASSBANK")

library(tidyverse)
library(tidymass)
library(plyr)

# data1 <-
#   read_msp_data_massbank(file = "MassBank_NIST.msp", threads = 5)
#
# massbank_nist_ms2 <-
#   convert_massbank2metid(data = data1,
#                          path = "NIST",
#                          source = "nist",
#                          threads = 5)
#
# load("NIST/massbank_ms2")
# massbank_nist_ms2 <- massbank_ms2
#
# colnames
# colnames
#
# idx1 <-
#   match(
#     massbank_nist_ms2@spectra.info$INCHI.ID,
#     hmdb_ms1@spectra.info$INCHI.ID,
#     incomparables = NA
#   )
```

```
#
# idx2 <-
#   match(
#     massbank_nist_ms2@spectra.info$INCHIKEY.ID,
#     hmdb_ms1@spectra.info$INCHIKEY.ID,
#     incomparables = NA
#   )
#
# idx3 <-
#   match(
#     massbank_nist_ms2@spectra.info$SMILES.ID,
#     hmdb_ms1@spectra.info$SMILES.ID,
#     incomparables = NA
#   )
#
# idx <-
#   data.frame(idx1, idx2, idx3) %>%
#   apply(1, function(x) {
#     x <- as.numeric(x)
#     x <- x[!is.na(x)]
#     if (length(x) == 0) {
#       return(NA)
#     }
#     return(x[1])
#   })
#
#
# cbind(
#   x = massbank_nist_ms2@spectra.info$Compound.name,
#   y = hmdb_ms1@spectra.info$Compound.name[idx]
# ) %>%
#   as.data.frame() %>%
#   dplyr::filter(!is.na(y))
#
# colnames
# colnames
#
# spectra.info <-
#  
#
# spectra.info <-
#   spectra.info %>%
#   dplyr::select(-c(CAS.ID, HMDB.ID, KEGG.ID))
#
# spectra.info <-
#   data.frame(spectra.info,
#             [idx, setdiff(colnames,
colnames(spectra.info))]) %>%
#   dplyr::select(-c(version, Create_date, Updated_date, secondary_accessions))
#
```

```
# # cbind(spectra.info$SMILES.ID,
# #       hmdb_ms1@spectra.info$SMILES.ID[idx])
#
#
# massbank_nist_ms2@spectra.info <-
#   spectra.info
#
# save(massbank_nist_ms2, file = "massbank_nist_ms2.rda")
#
# data2 <-
#   read_msp_data_massbank(file = "MassBank_RIKEN.msp", threads = 5)
#
# massbank_riken_ms2 <-
#   convert_massbank2metid(data = data2,
#                          threads = 5,
#                          source = "riken")
#
# new_spectra_info <-
# seq_along(massbank_riken_ms2@spectra.info$Comment) %>%
#   purrr::map(function(i){
#     # cat(i, " ")
#     x <- massbank_riken_ms2@spectra.info$Comment[i]
#     x <-
#     stringr::str_split(x, ";")[[1]] %>%
#       stringr::str_trim()
#     DB <- x[stringr::str_detect(x, "^DB")] %>%
#       stringr::str_replace("^DB#=", "") %>%
#       stringr::str_trim()
#     DB <- ifelse(length(DB) == 0, NA, DB)
#     origin <- x[stringr::str_detect(x, "^origin=")] %>%
#       stringr::str_replace("^origin=", "") %>%
#       stringr::str_trim()
#     origin <- ifelse(length(origin) == 0, NA, origin)
#     Comment_confidence <- x[stringr::str_detect(x, "^Annotation")] %>%
#       stringr::str_replace("^Annotation", "") %>%
#       stringr::str_trim()
#     Comment_confidence <- ifelse(length(Comment_confidence) == 0, NA,
Comment_confidence)
#     data.frame(DB, origin, Comment_confidence)
#   }) %>%
#   dplyr::bind_rows() %>%
#   as.data.frame()
#
# massbank_riken_ms2@spectra.info$Lab.ID <- new_spectra_info$DB
# massbank_riken_ms2@spectra.info$MASSBANK.ID <- new_spectra_info$DB
# massbank_riken_ms2@spectra.info$Submitter_team <- new_spectra_info$origin
# massbank_riken_ms2@spectra.info$Comment_confidence <-
new_spectra_info$Comment_confidence
#
# colnames
```

```
# colnames
#
# idx1 <-
#   match(massbank_riken_ms2@spectra.info$INCHI.ID,
#         hmdb_ms1@spectra.info$INCHI.ID,
#         incomparables = NA)
#
# idx2 <-
#   match(massbank_riken_ms2@spectra.info$INCHIKEY.ID,
#         hmdb_ms1@spectra.info$INCHIKEY.ID,
#         incomparables = NA)
#
# idx3 <-
#   match(massbank_riken_ms2@spectra.info$SMILES.ID,
#         hmdb_ms1@spectra.info$SMILES.ID,
#         incomparables = NA)
#
# idx <-
#   data.frame(idx1, idx2, idx3) %>%
#   apply(1, function(x) {
#     x <- as.numeric(x)
#     x <- x[!is.na(x)]
#     if (length(x) == 0) {
#       return(NA)
#     }
#     return(x[1])
#   })
#
#
# cbind(
#   x = massbank_riken_ms2@spectra.info$Compound.name,
#   y = hmdb_ms1@spectra.info$Compound.name[idx]
# ) %>%
#   as.data.frame() %>%
#   dplyr::filter(!is.na(y))
#
# colnames
# colnames
#
# spectra.info <-
#  
#
# spectra.info <-
#   spectra.info %>%
#   dplyr::select(-c(CAS.ID, HMDB.ID, KEGG.ID))
#
# spectra.info <-
#   data.frame(spectra.info,
#             [idx, setdiff(colnames,
colnames(spectra.info))]) %>%
```

```
#   dplyr::select(-c(version, Create_date, Updated_date, secondary_accessions))
#
#
# massbank_riken_ms2@spectra.info <-
#   spectra.info
#
# save(massbank_riken_ms2, file = "massbank_riken_ms2.rda")



# ####combine riken and nist
# load("massbank_riken_ms2.rda")
# load("massbank_nist_ms2.rda")
#
# intersect_compound_name <-
#   intersect(
#     massbank_nist_ms2@spectra.info$Compound.name,
#     massbank_riken_ms2@spectra.info$Compound.name
#   )
#
# intersect_compound_name[1]
# Lab.ID1 <-
#   %>%
#   dplyr::filter(Compound.name == intersect_compound_name[1]) %>%
#   pull(Lab.ID)
#
# Lab.ID2 <-
#   %>%
#   dplyr::filter(Compound.name == intersect_compound_name[1]) %>%
#   pull(Lab.ID)
#
# Lab.ID1
# Lab.ID2
#
# which(names(massbank_nist_ms2@spectra.data$Spectra.positive) == Lab.ID1[1])
# which(names(massbank_riken_ms2@spectra.data$Spectra.positive) == Lab.ID2[1])
#
# masstools::ms2_plot(
#   spectrum1 = massbank_nist_ms2@spectra.data$Spectra.positive[[1]][[1]],
#   spectrum2 = massbank_riken_ms2@spectra.data$Spectra.positive[[1]][[1]]
# )
#
# which(names(massbank_nist_ms2@spectra.data$Spectra.negative) == Lab.ID1[4])
# which(names(massbank_riken_ms2@spectra.data$Spectra.negative) == Lab.ID2[4])
#
#
# ##remove redundant spectra
# intersect_compound_name
# massbank_riken_ms2 <-
#   massbank_riken_ms2 %>%
```

```
#   dplyr::filter(!Compound.name %in% intersect_compound_name)
#
# massbank_ms2 <-
#   massbank_nist_ms2
#
# grep("[0-9]{1,5}\\.[0-9]{1,5}_[0-9]{1,5}\\.[0-9]{1,5}",
#      massbank_ms2@spectra.info$Compound.name,
#      value = FALSE)
#
# grep("[0-9]{1,5}\\.[0-9]{1,5}_[0-9]{1,5}\\.[0-9]{1,5}",
#      massbank_ms2@spectra.info$Compound.name,
#      value = TRUE)
#
#
massbank_ms2@spectra.info$Compound.name[!is.na(massbank_ms2@spectra.info$HMDB.ID)] <-
#
hmdb_ms1@spectra.info$Compound.name[match(massbank_ms2@spectra.info$HMDB[!is.na(massbank_ms2@spectra.info$HMDB.ID)],
#                                          hmdb_ms1@spectra.info$HMDB.ID)]
#
#
# rownames <- NULL
#
# save(massbank_ms2, file = "massbank_ms2.rda")
load("massbank_ms2.rda")
massbank_ms2

load("../HMDB/MS1/hmdb_ms1.rda")
load("../KEGG/kegg_ms1.rda")

intersect(colnames,
          colnames)

setdiff(colnames,
        colnames)

massbank_ms2 <-
  update_metid_database_info(
    database = massbank_ms2,
    ref_database = hmdb_ms1,
    by = c(
      "Compound.name",
      "INCHI.ID",
      "INCHIKEY.ID",
      "SMILES.ID",
      "CAS.ID",
      "HMDB.ID",
      "KEGG.ID",
      "IUPAC_name",
```

```
    "Traditional_IUPAC_name",
    "CHEMSPIDER.ID",
    "DRUGBANK.ID",
    "FOODB.ID",
    "PUBCHEM.ID",
    "CHEBI.ID",
    "BIOCYC.ID",
    "BIGG.ID",
    "WIKIPEDIA.ID",
    "METLIN.ID"
  ),
  combine_columns = c(
    "Compound.name",
    "INCHI.ID",
    "INCHIKEY.ID",
    "SMILES.ID",
    "Synonyms",
    "CAS.ID",
    "HMDB.ID",
    "KEGG.ID",
    "status",
    "Description",
    "monisotopic_molecular_weight",
    "IUPAC_name",
    "Traditional_IUPAC_name",
    "Kingdom",
    "Super_class",
    "Class",
    "Sub_class",
    "State",
    "Biospecimen_locations",
    "Cellular_locations",
    "Tissue_locations",
    "CHEMSPIDER.ID",
    "DRUGBANK.ID",
    "FOODB.ID",
    "PUBCHEM.ID",
    "CHEBI.ID",
    "BIOCYC.ID",
    "BIGG.ID",
    "WIKIPEDIA.ID",
    "METLIN.ID",
    "From_human"
  ),
  new_columns = c())

intersect(colnames,
        colnames)

setdiff(colnames,
```

```
        colnames)

source(here::here("R/3_utils.R"))

massbank_ms2 <-
  update_metid_database_info(
    database = massbank_ms2,
    ref_database = kegg_ms1,
    by = c(
      "Compound.name",
      "IUPAC_name",
      "Traditional_IUPAC_name",
      "CAS.ID",
      "SMILES.ID",
      "INCHI.ID",
      "INCHIKEY.ID",
      "FOODB.ID",
      "HMDB.ID",
      "PUBCHEM.ID",
      "CHEMSPIDER.ID",
      "KEGG.ID",
      "CHEBI.ID",
      "BIOCYC.ID",
      "WIKIPEDIA.ID",
      "DRUGBANK.ID",
      "BIGG.ID",
      "METLIN.ID"
    ),
    combine_columns = c(
      "Compound.name",
      "Synonyms",
      "IUPAC_name",
      "Traditional_IUPAC_name",
      "CAS.ID",
      "SMILES.ID",
      "INCHI.ID",
      "INCHIKEY.ID",
      "Kingdom",
      "Super_class",
      "Class",
      "Sub_class",
      "State",
      "FOODB.ID",
      "HMDB.ID",
      "PUBCHEM.ID",
      "CHEMSPIDER.ID",
      "KEGG.ID",
      "CHEBI.ID",
      "BIOCYC.ID",
      "WIKIPEDIA.ID",
```

```
    "status",
    "Biospecimen_locations",
    "Cellular_locations",
    "Tissue_locations",
    "DRUGBANK.ID",
    "BIGG.ID",
    "METLIN.ID",
    "From_human"
  ),
  new_columns = c(
    "CHEMBL.ID",
    "LIPIDMAPS.ID",
    "LIPIDBANK.ID",
    "From_drug",
    "KEGG_DRUG.ID"
  )
 )

massbank_ms2

load(here::here("other_files/source_system/source_system.rda"))

library(tidyverse)
library(tidyselect)
library(metid)

massbank_ms2 <-
 update_metid_database_source_system(
  database = massbank_ms2,
  source_system = source_system,
  by = c("CAS.ID", "HMDB.ID", "KEGG.ID"),
  prefer = "database"
 )

save(massbank_ms2, file = "massbank_ms2.rda")
```

**3. MoNA compound database**

```
###source
no_source()

masstools::setwd_project()
rm(list = ls())
load("other_files/HMDB/MS1/hmdb_ms1.rda")
load("other_files/KEGG/kegg_ms1.rda")
source("R/read_msp_data.R")
source("R/3_utils.R")
source("R/11_MONA.R")
```

```
setwd("other_files/MONA")

library(tidyverse)
library(tidymass)

# # #####CASMI2016
# # data1 <-
# #   read_msp_data(file = "MoNA-export-CASMI_2016.msp",
# #                 source = "mona",
# #                 threads = 5)
# #
# # mona_casmi2016_ms2 <-
# #   convert_mona2metid(data = data1,
# #                      path = "CASMI2016",
# #                      threads = 5)
# #
# # new_spectra_info <-
# #   mona_casmi2016_ms2@spectra.info$Comments %>%
# #   purrr::map(function(x) {
# #     x <-
# #       x %>%
# #       stringr::str_split(pattern = '\"') %>%
# #       `[[`(1) %>%
# #       stringr::str_trim()
# #     x <- x[x != ""]
# #     x <- x[x != " "]
# #     SMILES.ID =
# #       x[stringr::str_detect(x, "^SMILES")] %>% stringr::str_replace("SMILES=", "")
# #     SMILES.ID <- ifelse(length(SMILES.ID) == 0, NA, SMILES.ID)
# #     CAS.ID = x[stringr::str_detect(x, "^cas")] %>% stringr::str_replace("cas=", "")
# #     CAS.ID <- ifelse(length(CAS.ID) == 0, NA, CAS.ID)
# #     PUBCHEM.ID = x[stringr::str_detect(x, "^pubchem cid=")] %>%
stringr::str_replace("^pubchem cid=", "")
# #     PUBCHEM.ID <- ifelse(length(PUBCHEM.ID) == 0, NA, PUBCHEM.ID)
# #     CHEMSPIDER.ID = x[stringr::str_detect(x, "^chemspider=")] %>%
stringr::str_replace("^chemspider=", "")
# #     CHEMSPIDER.ID <-
# #       ifelse(length(CHEMSPIDER.ID) == 0, NA, CHEMSPIDER.ID)
# #     Author = x[stringr::str_detect(x, "^author=")] %>% stringr::str_replace("^author=", "")
# #     Author <- ifelse(length(Author) == 0, NA, Author)
# #     Comment_confidence <-
# #       x[stringr::str_detect(x, "^comment=CONFIDENCE")] %>%
stringr::str_replace("^comment=", "")
# #     Comment_confidence <-
# #       ifelse(length(Comment_confidence) == 0, NA, Comment_confidence)
# #     Submitter_team <-
# #       x[stringr::str_detect(x, "^submitter=")] %>% stringr::str_replace("^submitter=", "")
# #     Submitter_team <-
# #       ifelse(length(Submitter_team) == 0, NA, Submitter_team)
# #     data.frame(
```

```
##      SMILES.ID = SMILES.ID,
##      CAS.ID = CAS.ID,
##      PUBCHEM.ID = PUBCHEM.ID,
##      CHEMSPIDER.ID = CHEMSPIDER.ID,
##      Author = Author,
##      Comment_confidence = Comment_confidence,
##      Submitter_team = Submitter_team
##    )
##  }) %>%
##  dplyr::bind_rows() %>%
##  as.data.frame()
##
## colnames
## colnames(new_spectra_info)
## mona_casmi2016_ms2@spectra.info$SMILES.ID <-
##  new_spectra_info$SMILES.ID
## mona_casmi2016_ms2@spectra.info$CAS.ID <- new_spectra_info$CAS.ID
## mona_casmi2016_ms2@spectra.info$PUBCHEM.ID <-
##  new_spectra_info$PUBCHEM.ID
## mona_casmi2016_ms2@spectra.info$CHEMSPIDER.ID <-
##  new_spectra_info$CHEMSPIDER.ID
## mona_casmi2016_ms2@spectra.info$Author <- new_spectra_info$Author
## mona_casmi2016_ms2@spectra.info$Comment_confidence <-
##  new_spectra_info$Comment_confidence
## mona_casmi2016_ms2@spectra.info$Submitter_team <-
##  new_spectra_info$Submitter_team
##
## idx1 <-
##  match(
##   mona_casmi2016_ms2@spectra.info$INCHI.ID,
##   hmdb_ms1@spectra.info$INCHI.ID,
##   incomparables = NA
##  )
##
## idx2 <-
##  match(
##   mona_casmi2016_ms2@spectra.info$INCHIKEY.ID,
##   hmdb_ms1@spectra.info$INCHIKEY.ID,
##   incomparables = NA
##  )
##
## idx3 <-
##  match(
##   mona_casmi2016_ms2@spectra.info$CAS.ID,
##   hmdb_ms1@spectra.info$CAS.ID,
##   incomparables = NA
##  )
##
## idx4 <-
##  match(
```

```
##     mona_casmi2016_ms2@spectra.info$SMILES.ID,
##     hmdb_ms1@spectra.info$SMILES.ID,
##     incomparables = NA
##   )
##
## idx <-
##   data.frame(idx1, idx2, idx3, idx4) %>%
##   apply(1, function(x) {
##     x <- as.numeric(x)
##     x <- x[!is.na(x)]
##     if (length(x) == 0) {
##       return(NA)
##     }
##     return(x[1])
##   })
##
## colnames
## colnames
##
## spectra.info <-
##  
##
## spectra.info <-
##   spectra.info %>%
##   dplyr::select(-c(CAS.ID, HMDB.ID, KEGG.ID))
##
## spectra.info <-
##   data.frame(spectra.info,
##             [idx, setdiff(colnames,
colnames(spectra.info))]) %>%
##   dplyr::select(-c(version, Create_date, Updated_date, secondary_accessions))
##
##
## mona_casmi2016_ms2@spectra.info <-
##   spectra.info
##
## save(mona_casmi2016_ms2, file = "mona_casmi2016_ms2.rda")
##
##
##
##
##
##
## #####CASMI2012
## data1 <-
##   read_msp_data(file = "MoNA-export-CASMI_2012.msp",
##                 source = "mona",
##                 threads = 5)
##
## mona_casmi2012_ms2 <-
```

```
##   convert_mona2metid(data = data1,
##                      path = "CASMI2012",
##                      threads = 5)
##
## new_spectra_info <-
##   mona_casmi2012_ms2@spectra.info$Comments %>%
##   purrr::map(function(x) {
##     x <-
##       x %>%
##       stringr::str_split(pattern = '\"') %>%
##       `[[`(1) %>%
##       stringr::str_trim()
##     x <- x[x != ""]
##     x <- x[x != " "]
##     SMILES.ID =
##       x[stringr::str_detect(x, "^SMILES")] %>% stringr::str_replace("SMILES=", "")
##     SMILES.ID <- ifelse(length(SMILES.ID) == 0, NA, SMILES.ID)
##     CAS.ID = x[stringr::str_detect(x, "^cas")] %>% stringr::str_replace("cas=", "")
##     CAS.ID <- ifelse(length(CAS.ID) == 0, NA, CAS.ID)
##     PUBCHEM.ID = x[stringr::str_detect(x, "^pubchem cid=")] %>%
stringr::str_replace("^pubchem cid=", "")
##     PUBCHEM.ID <- ifelse(length(PUBCHEM.ID) == 0, NA, PUBCHEM.ID)
##     CHEMSPIDER.ID = x[stringr::str_detect(x, "^chemspider=")] %>%
stringr::str_replace("^chemspider=", "")
##     CHEMSPIDER.ID <-
##       ifelse(length(CHEMSPIDER.ID) == 0, NA, CHEMSPIDER.ID)
##     Author = x[stringr::str_detect(x, "^author=")] %>% stringr::str_replace("^author=", "")
##     Author <- ifelse(length(Author) == 0, NA, Author)
##     Comment_confidence <-
##       x[stringr::str_detect(x, "^comment=CONFIDENCE")] %>%
stringr::str_replace("^comment=", "")
##     Comment_confidence <-
##       ifelse(length(Comment_confidence) == 0, NA, Comment_confidence)
##     Submitter_team <-
##       x[stringr::str_detect(x, "^submitter=")] %>% stringr::str_replace("^submitter=", "")
##     Submitter_team <-
##       ifelse(length(Submitter_team) == 0, NA, Submitter_team)
##     data.frame(
##       SMILES.ID = SMILES.ID,
##       CAS.ID = CAS.ID,
##       PUBCHEM.ID = PUBCHEM.ID,
##       CHEMSPIDER.ID = CHEMSPIDER.ID,
##       Author = Author,
##       Comment_confidence = Comment_confidence,
##       Submitter_team = Submitter_team
##     )
##   }) %>%
##   dplyr::bind_rows() %>%
##   as.data.frame()
##
```

```
## colnames
## colnames(new_spectra_info)
## mona_casmi2012_ms2@spectra.info$SMILES.ID <-
##   new_spectra_info$SMILES.ID
## mona_casmi2012_ms2@spectra.info$CAS.ID <- new_spectra_info$CAS.ID
## mona_casmi2012_ms2@spectra.info$PUBCHEM.ID <-
##   new_spectra_info$PUBCHEM.ID
## mona_casmi2012_ms2@spectra.info$CHEMSPIDER.ID <-
##   new_spectra_info$CHEMSPIDER.ID
## mona_casmi2012_ms2@spectra.info$Author <- new_spectra_info$Author
## mona_casmi2012_ms2@spectra.info$Comment_confidence <-
##   new_spectra_info$Comment_confidence
## mona_casmi2012_ms2@spectra.info$Submitter_team <-
##   new_spectra_info$Submitter_team
##
## idx1 <-
##   match(
##     mona_casmi2012_ms2@spectra.info$INCHI.ID,
##     hmdb_ms1@spectra.info$INCHI.ID,
##     incomparables = NA
##   )
##
## idx2 <-
##   match(
##     mona_casmi2012_ms2@spectra.info$INCHIKEY.ID,
##     hmdb_ms1@spectra.info$INCHIKEY.ID,
##     incomparables = NA
##   )
##
## idx3 <-
##   match(
##     mona_casmi2012_ms2@spectra.info$CAS.ID,
##     hmdb_ms1@spectra.info$CAS.ID,
##     incomparables = NA
##   )
##
## idx4 <-
##   match(
##     mona_casmi2012_ms2@spectra.info$SMILES.ID,
##     hmdb_ms1@spectra.info$SMILES.ID,
##     incomparables = NA
##   )
##
## idx <-
##   data.frame(idx1, idx2, idx3, idx4) %>%
##   apply(1, function(x) {
##     x <- as.numeric(x)
##     x <- x[!is.na(x)]
##     if (length(x) == 0) {
##       return(NA)
```

```
##     }
##     return(x[1])
##   })
##
## colnames
## colnames
##
## spectra.info <-
##  
##
## spectra.info <-
##   spectra.info %>%
##   dplyr::select(-c(CAS.ID, HMDB.ID, KEGG.ID))
##
## spectra.info <-
##   data.frame(spectra.info,
##             [idx, setdiff(colnames,
colnames(spectra.info))]) %>%
##   dplyr::select(-c(version, Create_date, Updated_date, secondary_accessions))
##
##
## mona_casmi2012_ms2@spectra.info <-
##   spectra.info
##
## save(mona_casmi2012_ms2, file = "mona_casmi2012_ms2.rda")
##
##
##
##
##
##
##
##
## #####Fiehn HILIC
## data1 <-
##   read_msp_data(file = "MoNA-export-Fiehn_HILIC.msp",
##                 source = "mona",
##                 threads = 5)
##
## mona_fiehnhilic_ms2 <-
##   convert_mona2metid(data = data1,
##                      path = "Fiehn_HILIC",
##                      threads = 5)
##
## new_spectra_info <-
##   mona_fiehnhilic_ms2@spectra.info$Comments %>%
##   purrr::map(function(x) {
##     x <-
##       x %>%
##       stringr::str_split(pattern = '\"') %>%
```

```
##       `[[`(1) %>%
##       stringr::str_trim()
##     x <- x[x != ""]
##     x <- x[x != " "]
##     SMILES.ID =
##       x[stringr::str_detect(x, "^SMILES")] %>% stringr::str_replace("SMILES=", "")
##     SMILES.ID <- ifelse(length(SMILES.ID) == 0, NA, SMILES.ID)
##     CAS.ID = x[stringr::str_detect(x, "^cas")] %>% stringr::str_replace("cas=", "")
##     CAS.ID <- ifelse(length(CAS.ID) == 0, NA, CAS.ID)
##     PUBCHEM.ID = x[stringr::str_detect(x, "^pubchem cid=")] %>%
stringr::str_replace("^pubchem cid=", "")
##     PUBCHEM.ID <- ifelse(length(PUBCHEM.ID) == 0, NA, PUBCHEM.ID)
##     CHEMSPIDER.ID = x[stringr::str_detect(x, "^chemspider=")] %>%
stringr::str_replace("^chemspider=", "")
##     CHEMSPIDER.ID <-
##       ifelse(length(CHEMSPIDER.ID) == 0, NA, CHEMSPIDER.ID)
##     Author = x[stringr::str_detect(x, "^author=")] %>% stringr::str_replace("^author=", "")
##     Author <- ifelse(length(Author) == 0, NA, Author)
##     Comment_confidence <-
##       x[stringr::str_detect(x, "^comment=CONFIDENCE")] %>%
stringr::str_replace("^comment=", "")
##     Comment_confidence <-
##       ifelse(length(Comment_confidence) == 0, NA, Comment_confidence)
##     Submitter_team <-
##       x[stringr::str_detect(x, "^submitter=")] %>% stringr::str_replace("^submitter=", "")
##     Submitter_team <-
##       ifelse(length(Submitter_team) == 0, NA, Submitter_team)
##     data.frame(
##       SMILES.ID = SMILES.ID,
##       CAS.ID = CAS.ID,
##       PUBCHEM.ID = PUBCHEM.ID,
##       CHEMSPIDER.ID = CHEMSPIDER.ID,
##       Author = Author,
##       Comment_confidence = Comment_confidence,
##       Submitter_team = Submitter_team
##     )
##   }) %>%
##   dplyr::bind_rows() %>%
##   as.data.frame()
##
## colnames
## colnames(new_spectra_info)
## mona_fiehnhilic_ms2@spectra.info$SMILES.ID <-
##   new_spectra_info$SMILES.ID
## mona_fiehnhilic_ms2@spectra.info$CAS.ID <- new_spectra_info$CAS.ID
## mona_fiehnhilic_ms2@spectra.info$PUBCHEM.ID <-
##   new_spectra_info$PUBCHEM.ID
## mona_fiehnhilic_ms2@spectra.info$CHEMSPIDER.ID <-
##   new_spectra_info$CHEMSPIDER.ID
## mona_fiehnhilic_ms2@spectra.info$Author <- new_spectra_info$Author
```

```
## mona_fiehnhilic_ms2@spectra.info$Comment_confidence <-
##   new_spectra_info$Comment_confidence
## mona_fiehnhilic_ms2@spectra.info$Submitter_team <-
##   new_spectra_info$Submitter_team
##
## idx1 <-
##   match(
##     mona_fiehnhilic_ms2@spectra.info$INCHI.ID,
##     hmdb_ms1@spectra.info$INCHI.ID,
##     incomparables = NA
##   )
##
## idx2 <-
##   match(
##     mona_fiehnhilic_ms2@spectra.info$INCHIKEY.ID,
##     hmdb_ms1@spectra.info$INCHIKEY.ID,
##     incomparables = NA
##   )
##
## idx3 <-
##   match(
##     mona_fiehnhilic_ms2@spectra.info$CAS.ID,
##     hmdb_ms1@spectra.info$CAS.ID,
##     incomparables = NA
##   )
##
## idx4 <-
##   match(
##     mona_fiehnhilic_ms2@spectra.info$SMILES.ID,
##     hmdb_ms1@spectra.info$SMILES.ID,
##     incomparables = NA
##   )
##
## idx <-
##   data.frame(idx1, idx2, idx3, idx4) %>%
##   apply(1, function(x) {
##     x <- as.numeric(x)
##     x <- x[!is.na(x)]
##     if (length(x) == 0) {
##       return(NA)
##     }
##     return(x[1])
##   })
##
## colnames
## colnames
##
## spectra.info <-
##  
##
```

```
# # spectra.info <-
# #   spectra.info %>%
# #   dplyr::select(-c(CAS.ID, HMDB.ID, KEGG.ID))
# #
# # spectra.info <-
# #   data.frame(spectra.info,
# #             [idx, setdiff(colnames,
colnames(spectra.info))]) %>%
# #   dplyr::select(-c(version, Create_date, Updated_date, secondary_accessions))
# #
# # mona_fiehnhilic_ms2@spectra.info <-
# #   spectra.info
# #
# # save(mona_fiehnhilic_ms2, file = "mona_fiehnhilic_ms2.rda")
# #
# #
# #
# #
# #
# #
# #
# #
# # #####EMBL-MCF
# # data1 <-
# #   read_msp_data(file = "MoNA-export-EMBL-MCF.msp",
# #                 source = "mona",
# #                 threads = 5)
# #
# # mona_emblmcf_ms2 <-
# #   convert_mona2metid(data = data1,
# #                      path = "EMBL-MCF",
# #                      threads = 5)
# #
# # new_spectra_info <-
# #   mona_emblmcf_ms2@spectra.info$Comments %>%
# #   purrr::map(function(x) {
# #     x <-
# #       x %>%
# #       stringr::str_split(pattern = '\"') %>%
# #       `[[`(1) %>%
# #       stringr::str_trim()
# #     x <- x[x != ""]
# #     x <- x[x != " "]
# #     SMILES.ID =
# #       x[stringr::str_detect(x, "^SMILES")] %>% stringr::str_replace("SMILES=", "")
# #     SMILES.ID <- ifelse(length(SMILES.ID) == 0, NA, SMILES.ID)
# #     CAS.ID = x[stringr::str_detect(x, "^cas")] %>% stringr::str_replace("cas=", "")
# #     CAS.ID <- ifelse(length(CAS.ID) == 0, NA, CAS.ID)
# #     PUBCHEM.ID = x[stringr::str_detect(x, "^pubchem cid=")] %>%
stringr::str_replace("^pubchem cid=", "")
```

```
##     PUBCHEM.ID <- ifelse(length(PUBCHEM.ID) == 0, NA, PUBCHEM.ID)
##     CHEMSPIDER.ID = x[stringr::str_detect(x, "^chemspider=")] %>%
stringr::str_replace("^chemspider=", "")
##     CHEMSPIDER.ID <-
##       ifelse(length(CHEMSPIDER.ID) == 0, NA, CHEMSPIDER.ID)
##     Author = x[stringr::str_detect(x, "^author=")] %>% stringr::str_replace("^author=", "")
##     Author <- ifelse(length(Author) == 0, NA, Author)
##     Comment_confidence <-
##       x[stringr::str_detect(x, "^comment=CONFIDENCE")] %>%
stringr::str_replace("^comment=", "")
##     Comment_confidence <-
##       ifelse(length(Comment_confidence) == 0, NA, Comment_confidence)
##     Submitter_team <-
##       x[stringr::str_detect(x, "^submitter=")] %>% stringr::str_replace("^submitter=", "")
##     Submitter_team <-
##       ifelse(length(Submitter_team) == 0, NA, Submitter_team)
##     data.frame(
##       SMILES.ID = SMILES.ID,
##       CAS.ID = CAS.ID,
##       PUBCHEM.ID = PUBCHEM.ID,
##       CHEMSPIDER.ID = CHEMSPIDER.ID,
##       Author = Author,
##       Comment_confidence = Comment_confidence,
##       Submitter_team = Submitter_team
##     )
##   }) %>%
##   dplyr::bind_rows() %>%
##   as.data.frame()
##
## colnames
## colnames(new_spectra_info)
## mona_emblmcf_ms2@spectra.info$SMILES.ID <-
##   new_spectra_info$SMILES.ID
## mona_emblmcf_ms2@spectra.info$CAS.ID <-
##   new_spectra_info$CAS.ID
## mona_emblmcf_ms2@spectra.info$PUBCHEM.ID <-
##   new_spectra_info$PUBCHEM.ID
## mona_emblmcf_ms2@spectra.info$CHEMSPIDER.ID <-
##   new_spectra_info$CHEMSPIDER.ID
## mona_emblmcf_ms2@spectra.info$Author <-
##   new_spectra_info$Author
## mona_emblmcf_ms2@spectra.info$Comment_confidence <-
##   new_spectra_info$Comment_confidence
## mona_emblmcf_ms2@spectra.info$Submitter_team <-
##   new_spectra_info$Submitter_team
##
## idx1 <-
##   match(
##     mona_emblmcf_ms2@spectra.info$INCHI.ID,
##     hmdb_ms1@spectra.info$INCHI.ID,
```

```
##     incomparables = NA
##   )
##
## idx2 <-
##   match(
##     mona_emblmcf_ms2@spectra.info$INCHIKEY.ID,
##     hmdb_ms1@spectra.info$INCHIKEY.ID,
##     incomparables = NA
##   )
##
## idx3 <-
##   match(
##     mona_emblmcf_ms2@spectra.info$CAS.ID,
##     hmdb_ms1@spectra.info$CAS.ID,
##     incomparables = NA
##   )
##
## idx4 <-
##   match(
##     mona_emblmcf_ms2@spectra.info$SMILES.ID,
##     hmdb_ms1@spectra.info$SMILES.ID,
##     incomparables = NA
##   )
##
## idx <-
##   data.frame(idx1, idx2, idx3, idx4) %>%
##   apply(1, function(x) {
##     x <- as.numeric(x)
##     x <- x[!is.na(x)]
##     if (length(x) == 0) {
##       return(NA)
##     }
##     return(x[1])
##   })
##
## colnames
## colnames
##
## spectra.info <-
##  
##
## spectra.info <-
##   spectra.info %>%
##   dplyr::select(-c(CAS.ID, HMDB.ID, KEGG.ID))
##
## spectra.info <-
##   data.frame(spectra.info,
##             [idx, setdiff(colnames,
colnames(spectra.info))]) %>%
##   dplyr::select(-c(version, Create_date, Updated_date, secondary_accessions))
```

```
# #
# # mona_emblmcf_ms2@spectra.info <-
# #   spectra.info
# #
# # save(mona_emblmcf_ms2, file = "mona_emblmcf_ms2.rda")
#
#
# load("mona_casmi2012_ms2.rda")
# load("mona_casmi2016_ms2.rda")
# load("mona_emblmcf_ms2.rda")
# load("mona_fiehnhilic_ms2.rda")
#
#
# mona_casmi2012_ms2
# mona_casmi2016_ms2
#
# intersect(
#   mona_casmi2012_ms2@spectra.info$Compound.name,
#   mona_casmi2016_ms2@spectra.info$Compound.name
# )
#
# colnames ==
# colnames
#
# mona_casmi2012_ms2@spectra.info$Lab.ID
# mona_casmi2016_ms2@spectra.info$Lab.ID
#
# names(mona_casmi2012_ms2@spectra.data$Spectra.positive)
# names(mona_casmi2012_ms2@spectra.data$Spectra.negative)
#
# mona_ms2 <-
#   mona_casmi2012_ms2
#
# mona_casmi2012_ms2@spectra.info$Lab.ID
# mona_casmi2016_ms2@spectra.info$Lab.ID
# mona_emblmcf_ms2@spectra.info$Lab.ID
# mona_fiehnhilic_ms2@spectra.info$Lab.ID
#
# mona_ms2@spectra.info <-
#   rbind(,
#        ,
#        ,
#        )
#
# mona_ms2@spectra.data$Spectra.positive <-
#   c(mona_casmi2012_ms2@spectra.data$Spectra.positive,
#     mona_casmi2016_ms2@spectra.data$Spectra.positive,
#     mona_emblmcf_ms2@spectra.data$Spectra.positive,
#     mona_fiehnhilic_ms2@spectra.data$Spectra.positive)
#
```

```
# mona_ms2@spectra.data$Spectra.negative <-
#   c(mona_casmi2012_ms2@spectra.data$Spectra.negative,
#     mona_casmi2016_ms2@spectra.data$Spectra.negative,
#     mona_emblmcf_ms2@spectra.data$Spectra.negative,
#     mona_fiehnhilic_ms2@spectra.data$Spectra.negative)
#
# ####remove labeled compounds
# mona_ms2@spectra.info$Submitter_team
#
# mona_ms2@spectra.info$Compound.name
#
# grep("\\&", mona_ms2@spectra.info$Compound.name, value = TRUE)
# mona_ms2@spectra.info$INCHIKEY.ID[grep("\\&",
mona_ms2@spectra.info$Compound.name, value = FALSE)]
#
# mona_ms2@spectra.info$Compound.name <-
# mona_ms2@spectra.info$Compound.name %>%
#   stringr::str_replace_all("&\\#39\\;", "")
#
# save(mona_ms2, file = "mona_ms2.rda")
load("mona_ms2.rda")

load("../HMDB/MS1/hmdb_ms1.rda")
load("../KEGG/kegg_ms1.rda")

intersect(colnames,
          colnames)

setdiff(colnames,
        colnames)

mona_ms2 <-
  update_metid_database_info(
    database = mona_ms2,
    ref_database = hmdb_ms1,
    by = c(
      "Compound.name",
      "CAS.ID",
      "HMDB.ID",
      "KEGG.ID",
      "METLIN.ID"
    ),
    combine_columns = c(
      "Compound.name",
      "CAS.ID",
      "HMDB.ID",
      "KEGG.ID",
      "METLIN.ID"
    ),
    new_columns = c())
```

```
intersect(colnames,
          colnames)

setdiff(colnames,
        colnames)

source(here::here("R/3_utils.R"))

mona_ms2 <-
  update_metid_database_info(
    database = mona_ms2,
    ref_database = kegg_ms1,
    by = c(
      "Compound.name",
      "IUPAC_name",
      "Traditional_IUPAC_name",
      "CAS.ID",
      "SMILES.ID",
      "INCHI.ID",
      "INCHIKEY.ID",
      "FOODB.ID",
      "HMDB.ID",
      "PUBCHEM.ID",
      "CHEMSPIDER.ID",
      "KEGG.ID",
      "CHEBI.ID",
      "BIOCYC.ID",
      "WIKIPEDIA.ID",
      "DRUGBANK.ID",
      "BIGG.ID",
      "METLIN.ID"
    ),
    combine_columns = c(
      "Compound.name",
      "Synonyms",
      "IUPAC_name",
      "Traditional_IUPAC_name",
      "CAS.ID",
      "SMILES.ID",
      "INCHI.ID",
      "INCHIKEY.ID",
      "Kingdom",
      "Super_class",
      "Class",
      "Sub_class",
      "State",
      "FOODB.ID",
      "HMDB.ID",
      "PUBCHEM.ID",
```

```
    "CHEMSPIDER.ID",
    "KEGG.ID",
    "CHEBI.ID",
    "BIOCYC.ID",
    "WIKIPEDIA.ID",
    "status",
    "Biospecimen_locations",
    "Cellular_locations",
    "Tissue_locations",
    "DRUGBANK.ID",
    "BIGG.ID",
    "METLIN.ID",
    "From_human"
   ),
   new_columns = c(
    "CHEMBL.ID",
    "LIPIDMAPS.ID",
    "LIPIDBANK.ID",
    "From_drug",
    "KEGG_DRUG.ID"
   )
  )

load(here::here("other_files/source_system/source_system.rda"))

library(tidyverse)
library(tidyselect)
library(metid)

mona_ms2 <-
  update_metid_database_source_system(
   database = mona_ms2,
   source_system = source_system,
   by = c("CAS.ID", "HMDB.ID", "KEGG.ID"),
   prefer = "database"
  )

save(mona_ms2, file = "mona_ms2.rda")
```

**4. KEGG pathway database**

```
library(massdatabase)
pathway_info <-
  request_kegg_pathway_info(organism = "hsa")
head(pathway_info)
pathway <-
  request_kegg_pathway(pathway_id = "hsa00010")
pathway
download_kegg_pathway(path = "kegg_human_pathway",
```

```
                      sleep = 1,
                      organism = "hsa")
data <-
  read_kegg_pathway(path = "kegg_human_pathway")
class(data)
 kegg_human_pathway <-
 convert_kegg2metpath(data = data,
                      path = "kegg_human_pathway",
                     threads = 3)
```
